# Mutation effect estimation on protein-protein interactions using deep contextualized representation learning

**DOI:** 10.1101/2019.12.15.876953

**Authors:** Guangyu Zhou, Muhao Chen, Chelsea J.-T. Ju, Zheng Wang, Jyun-Yu Jiang, Wei Wang

## Abstract

The functional impact of protein mutations is reflected on the alteration of conformation and thermodynamics of protein-protein interactions (PPIs). Quantifying the changes of two interacting proteins upon mutations are commonly carried out by computational approaches. Hence, extensive research efforts have been put to the extraction of energetic or structural features on proteins, followed by statistical learning methods to estimate the effects of mutations to PPI properties. Nonetheless, such features require extensive human labors and expert knowledge to obtain, and have limited abilities to reflect point mutations. We present an end-to-end deep learning framework, `MuPIPR`, to estimate the effects of mutations on PPIs. `MuPIPR` incorporates a contextualized representation mechanism of amino acids to propagate the effects of a point mutation to surrounding amino acid representations, therefore amplifying the subtle change in a long protein sequence. On top of that, `MuPIPR` leverages a Siamese residual recurrent convolutional neural encoder to encode a wildtype protein pair and its mutation pair. Multiple-layer perceptron regressors are applied to the protein pair representations to predict the quantifiable changes of PPI properties upon mutations. Experimental evaluations show that `MuPIPR` outperforms various state-of-the-art systems on the change of binding affinity prediction and the buried surface area prediction. The software implementation is available at https://github.com/guangyu-zhou/MuPIPR

## 1 Introduction

Protein-protein interactions (PPIs) govern a wide range of biological mechanisms ranging from metabolic and signalling pathways, cellular processes, and immune system [15, 44]. Mutations in proteins can affect protein folding and stability [1, 34, 35]; consequently, these mutations alter the kinetic and thermodynamics of PPIs [21, 26]. Such mutations can either be selectively advantageous to the organism through evolution [43], or be deleterious and causing a disease phenotype [24]. Understanding the effects of these mutations, specifically the conformation and thermodynamic changes of the interacting proteins, is vital to various biomedical applications, including disease-associated mutation analyses [46], drug design [17], and therapeutic intervention [14].

The quantitative measures of different aspects of PPIs are often determined experimentally through biophysical techniques, including but not limited to, isothermal titration calorimetry for measuring the binding affinity [27], and BN-PAGE for the stability of the protein complex [40]. However, these *in vivo* or *in vitro* techniques are laborious and expensive due to the need of purifying each protein. In addition, estimating the mutation effects requires the physical presence of both wild-type and mutated proteins, while the interaction involving a mutated protein is not always available.

To enable large scale studies of the mutation effects of PPIs, significant efforts have been made to computationally estimating the changes of PPIs properties upon mutations. One of these estimates, the change of binding affinity (*∆∆G*), has been widely investigated. The binding affinity (*∆G*) measures the strength of the interaction between two single biomolecules, and is reported by the equilibrium dissociation constant. Here, the basis of a biomolecule can be a protein or a subunit of a protein. Earlier methods estimating the *∆∆G* derive empirical linear functions based on physical energies [16] or known protein structures [11, 33]. Instead of training parameters on existing data, these methods require a set of pre-determined coefficients for the linear models. Hence, they suffer from poor generalizability and lead to low accuracy. With the increasing availability of large mutation databases, statistical learning algorithms have been proposed to capture the relations between a variety of energetic or structural features and the binding affinity of two biomolecules [13, 29, 43, 51]. Nevertheless, features used in these methods are hand-crafted, requiring extensive human labors and expert knowledge.

Recently, deep learning methods show the potential of automatically extracting comprehensive features from protein sequences, and gain unprecedented success in PPI tasks. Corresponding methods, including DNN-PPI [28], DPPI [18], and PIPR [8] employ various neural sequence pair models to predict PPI information based on protein sequences. In the application of predicting the structural properties of a protein complex, NetSurfp-2.0 [25] employs a neural sequence model to predict the solvent accessible surface area (ASA) of a protein. A more pronounced feature to describe the PPI property is the buried surface area (BSA), which measures the size of the interface in a protein-protein complex. Estimating the change of BSA upon mutation provides insight to the potential deleterious mutations buried inside an interacting protein pair, which is crucial for understanding diseases [7]. However, none of the existing methods can directly estimate the BSA score, or the change in BSA upon mutation. BSA is often computed from the ASA scores of individual proteins and the protein complex [13].

To capture the features of the raw protein sequence from scratch, neural sequence models require a mechanism to characterize and represent the amino acids in a protein sequence. Such mechanisms deploy fixed representations, e.g. one-hot vectors [28], physicochemical property-aware encoding [8] or static amino acid embeddings [8]. However, these representations face several indispensable problems for PPI property predictions upon mutations. First, these representation mechanisms fall short of reflecting mutations, inasmuch as point mutations will only lead to subtle differences in the corresponding amino acid representations of a long protein sequence. Second, as these mechanisms independently characterize each amino acid, they do not consider the contextual information of surrounding amino acids and do not highlight those that are critical to PPIs. Moreover, the aforementioned methods fail to deploy a learning architecture dedicated to modeling the change between a wide-type protein pair and its mutant counterparts.

In this paper, we introduce a comprehensive learning framework, `MuPIPR` (<u>Mu</u>tation Effects in <u>P</u>rotein-protein Interaction <u>PR</u>ediction Using Contextualized Representations), to estimate the changes of quantifiable PPI properties upon amino acid mutations. `MuPIPR` incorporates a contextualized representation learning mechanism of amino acids, which seeks to sensitively capture mutations. In particular, `MuPIPR` pre-trains a multi-layer bidirectional long short-term memory (LSTM) [20] language model on a collection of protein sequences, and alters the representation of each amino acid based on the surrounding context captured by the language model. The benefits of this representation learning mechanism are two-fold: (i) it automatically extracts more refined amino-acid-level features that are differentiated between different contexts of the proteins; (ii) it propagates the mutation effects to the representations of surrounding amino acids, therefore amplifying the subtle signal of each mutation in a long protein sequence. On top of the contextualized animo acid representations, a deep neural learning architecture is carefully designed to subsequently estimate the quantifiable PPI property changes between a wild-type pair and a mutant pair of proteins. The architecture features an end-to-end learning of two stages: (i) a 4-fold Siamese residual recurrent convolutional neural network (RCNN) encoder characterizes the two protein pairs based on their contextualized amino acid representations, which seeks to seize the differences of their latent features; (ii) a main multi-layer perceptron (MLP) regressor is stacked to the encoder to estimate the property change between the two protein pairs, with two auxiliary regressors to enhance the estimation by jointly estimating the individual PPI properties of each protein pair. In practice, `MuPIPR` demonstrates the benefits of alleviating the need of hand-crafted features, and generalizes well to tasks of estimating the changes of different PPI properties upon mutations. The evaluation on the protein binding affinity change estimation task and protein buried surface area change estimation task on three benchmark datasets shows that `MuPIPR` significantly outperforms state-of-the-art methods on both tasks. Detailed ablation study on `MuPIPR` also provides insightful understanding of the effectiveness of each model component.

## 2 Methods

In this section, we present the detailed design of the proposed framework `MuPIPR` to address two regression tasks in PPI. Figure 1 illustrates the architecture of `MuPIPR` with three components: (1) a contextualized amino acid representation mechanism, (2) a protein sequence level Siamese encoder by leveraging the residual recurrent convolutional neural network (RCNN), (3) multiple-layer perceptron regressors for estimating quantifiable property changes in PPIs upon mutations.

**Fig. 1:**
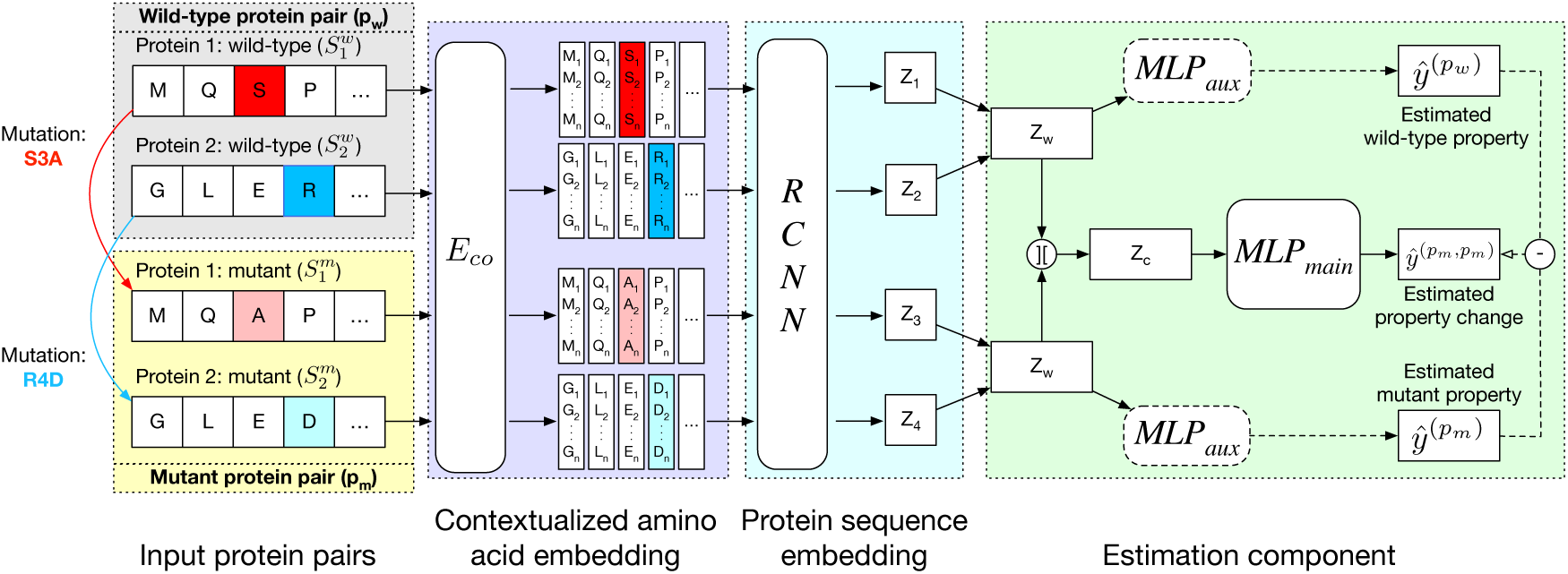
Architecture of `MuPIPR`.

### 2.1 Preliminary

We represent a protein as a sequence of amino acid residues *S* = (*r*_1_, *r*_2_, …, *r*_*N*_), where each *r*_*i*_ is an amino acid residue. Let *I* = {(*p*_*w*_, *p*_*m*_)} be a set of doublets, where each doublet contains a wild-type protein pair 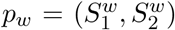 and its corresponding mutant protein pair 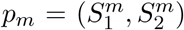. The mutant pair contains at least one mutant of the proteins from the wild-type pair, such that 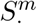 could be the mutant form of the 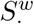. Our goal is to estimate the change of quantifiable PPI properties between a wild-type pair and its mutant pair. Such changes can be that of binding affinity, buried surface area, or other quantifiable properties.

### 2.2 Protein sequence encoding with contextualized representations

Primary sequence is the fundamental information to describe a protein. Apprehending the sequence information serves as the basis of estimating the effects caused by protein mutations. To better characterize the sequence information and the mutation, `MuPIPR` adopts two levels of encoding processes respectively on individual amino acids and on the sequence.

#### Contextualized amino acid embedding

Given an amino acid in a protein sequence, we seek to generate an adaptive representation according to its surrounding amino acids (referred as the context). The contextualized amino acid embedding produces a sequence of vector representations for the amino acid residues, 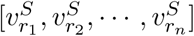, where each vector 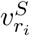 is the representation of residue *r*_*i*_ that is specific to the context of protein *S*. The detailed framework of training the contextualized amino acid embedding (*E*_*co*_) is shown in Figure 2a.

**Fig. 2:**
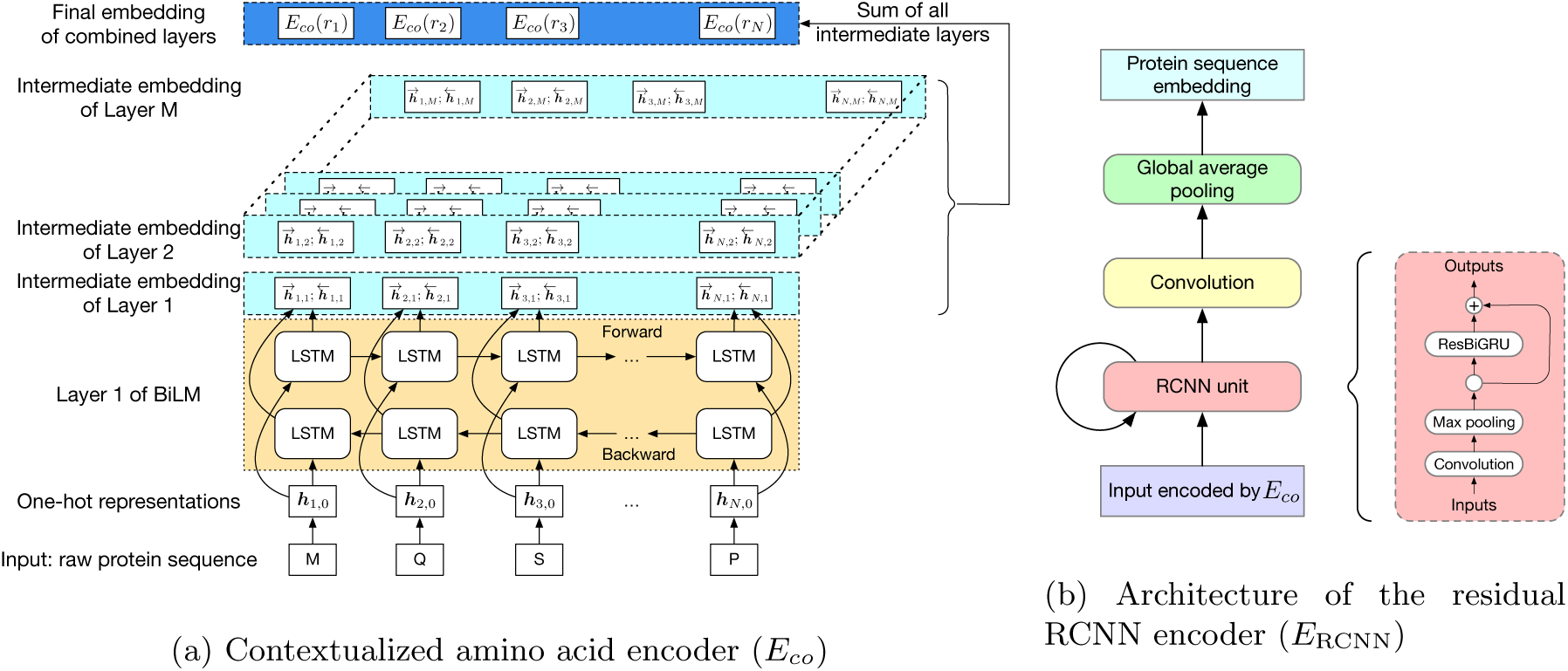
The detailed framework of the two encoders.

Compared to the static representations of the amino acid such as the one-hot encoding and co-occurrence based embeddings [8], the contextualized embedding seeks to incorporate the information of neighboring amino acids. Inspired by the recent success of ELMo [42] for word representations under different linguistic contexts, we obtain the contextualized representations of amino acids by leveraging a pre-trained bidirectional language model (BiLM). The BiLM is crucial to capturing the context information of a given amino acid in a sequence.

The forward language model computes the sequence probability of residue *r*_*i*_ given the context (*r*_1_, *r*_2_, …, *r*_*i*−1_):

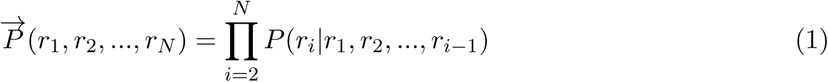

The backward language model computes the sequence probability in the reverse order. Given the later context, (*r*_*i*+1_, *r*_*i*+2_…, *r*_*N*_), it predicts the previous residue as:

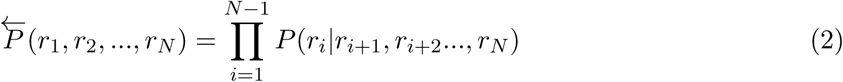

The language model of each direction is implemented with *M* stacked layers of long short-term memory (LSTM) models [20]. The LSTM layers of both directions output intermediate embedding vectors (i.e., hidden state vectors) 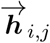 or 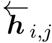 based on the context for the forward and backward language models respectively, where *j* = 1, …*M*. These intermediate embedding vectors from different layers represent different levels of contextual information. The vectors from higher-level layers capture the information of more long-term contexts, while lower-level vectors extract more fine-grained information of the neighboring amino acids. Specially, the output of the LSTM’s top layer, 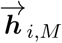 or 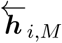, is passed through a Softmax layer to predict the next or previous residue *r*_*i*+1_ or *r*_*i*−1_.

By combining the above two language models, the objective of the BiLM is defined as follows:

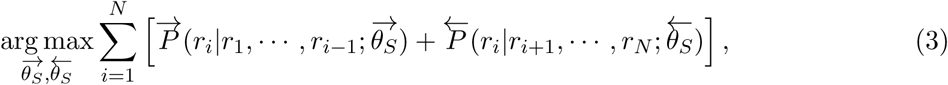

where 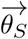 and 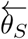 are the learnable parameters of either direction of LSTM.

We pre-train our BiLM on a selected set of raw protein sequences from the STRING protein database [48], which is described in detail in the Datasets Section. Then the pre-trained BiLM can be applied on each protein sequence of up to *N* amino acid residues. For each amino acid *r*_*i*_, the *M* -layer BiLM computes a total of 2*M* + 1 embeddings, denoted as *E*(*r*_*i*_) = {***h***_*i,j*_|*j* = 0, …, *M*}. Here, ***h***_*i*,0_ is a trainable single-layer affine projection [30] on one-hot vectors of amino acids and 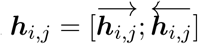 are the intermediate results for each layer of the BiLM. Different layers of the stacked BiLM capture different widths of the neighbouring contexts of the amino acid in the sequence. Therefore, for each amino acid, we aggregate the representations of different BiLM layers to obtain its contextualized representation: 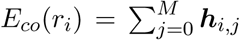. Then the encoding for *S* = (*r*_1_, *r*_2_, …, *r*_*N*_) is produced as a sequence of contextualized amino acid embedding vectors: *E*_*co*_(*S*) = [*E*_*co*_(*r*_1_), *E*_*co*_(*r*_2_), …, *E*_*co*_(*r*_*N*_)].

#### Protein sequence encoding

Upon obtaining the embeddings of amino acids for the protein sequence, we employ a deep 4-fold Siamese architecture of neural network to capture latent semantic features of the protein sequence doublets. Following PIPR [8], we choose to build a sequence level encoder based on the residual recurrent convolutional neural network (RCNN), due to its benefits with sequential and multi-granular feature aggregation for the protein sequence. `MuPIPR` can be easily adapted to use other sequence encoding techniques such as convolutional neural network (CNN) [18, 28] or self-attentive encoders [9, 49].

As depicted in Figure 2b, the residual RCNN sequence encoding process *E*_RCNN_ starts with an iterative process of the residual RCNN unit, consisting of two modules: a convolution module and a recurrent module. The convolution module serves as the initial encoder to extract local features from the input, followed by a recurrent module. Then the output of the recurrent module is used as the input of the next convolution module. This iterative process helps to generate and aggregate features while maintaining the contextualized features across different layers.

The convolution module contains a convolution layer with pooling. Let *X* be the input sequence of vectors. The convolution layer (Conv) applies a weight-sharing kernel of size *k* to generate a latent vector ***h***_*i*_ from each *k*-gram *X*_*i:i*+*k*_ of *X*. By sliding through the whole sequence, the convolution layer produces a sequence of the latent vectors: ***H***^(1)^ = [***h***_1_, ***h***_2_, …, ***h***_*|X|*+1−*k*_]. Then the *p*-max-pooling applies on every *p*-strides of the sequence (i.e. non-overlapping subsequences of length *p*). We use the max-pooling mechanism to preserve the most significant features within each stride. The output of this module is summarized as below:

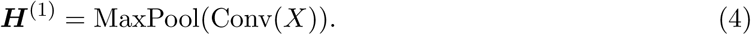

The recurrent module consists of the bidirectional gated recurrent units (BiGRU). Note that the gated recurrent units (GRU) is an alternative to LSTM without extra parameters on memory gates. Typically GRU performs better than LSTM on small data [10]. Consider that there are much fewer data for protein pairs with mutations than the raw sequence data for pre-training the BiLM, we use GRU instead of LSTM in this module. We also add the residual mechanism, which is designed to improve the learning of non-linear neural layers [19]. Given an input vector ***v***, the recurrent module with residual is defined as: 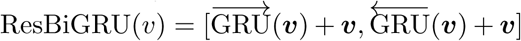. A residual RCNN unit stacks two modules:

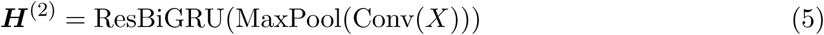

After a chain of residual RCNN units, the output of the last iteration 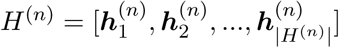 is passed through a final convolution layer with global average pooling [32]. This produces the final sequence embedding:

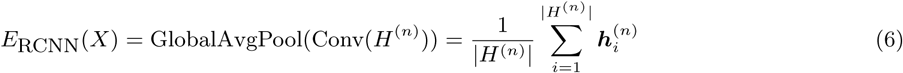

### 2.3 Design and learning objective of MuPIPR

#### Learning architecture

In order to estimate the changes of PPI properties upon mutations, our model needs to handle the inputs of four sequences of a doublet (*p*_*w*_, *p*_*m*_) at a time, i.e, two from a wild-type pair *p*_*w*_ and the other two from a mutant pair *p*_*w*_. Therefore, we deploy a 4-fold Siamese architecture to capture the mutual interaction between *p*_*w*_ and *p*_*m*_.

Given two protein pairs 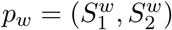 and 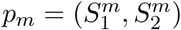, where 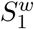, 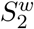 and 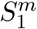, 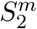 are the wild-type proteins and their mutant respectively, the pre-trained contextualized model is applied to obtain the amino acid level representations for each sequence: 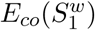, 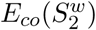, 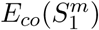, 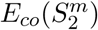. The amino acid embeddings of these four sequences are then served as inputs to one RCNN encoder, which yields 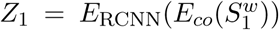, 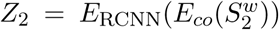, 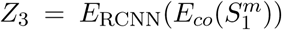, and 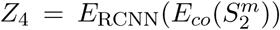. The encodings of two proteins in each pair are then combined by the concatenation of their element-wise product (*⊙*) and their absolute element-wise differences: *Z*_*w*_ = [*Z*_1_ ⊙ *Z*_2_; |*Z*_1_ −*Z*_2_|], and *Z*_*m*_ = [*Z*_3_ ⊙ *Z*_4_; |*Z*_3_ −*Z*_4_|], where *Z*_*w*_ and *Z*_*m*_ refer to the final encoding of the wild-type pair and the mutant pair, respectively. Intuitively, we use the integration technique defined for *Z*_*w*_ and *Z*_*m*_ to apprehend the mutual influence of a pair of protein sequences, as this is a widely used technique for symmetric pairwise comparison in neural sequence pair modeling. Afterwards, *Z*_*c*_ is obtained by the ordered concatenation of *Z*_*w*_ and *Z*_*m*_, i.e., *Z*_*c*_ = [*Z*_*w*_; *Z*_*m*_], as we specify the second pair to be the mutant one.

#### Primary learning objective

The property change upon mutation is estimated with a multi-layer perceptron (MLP) based regressor. Particularly, a MLP with leaky ReLU [36] is stacked on *Z*_*c*_, and outputs a scalar to estimate the PPI property change. The main learning objective is to minimize the following mean squared loss:

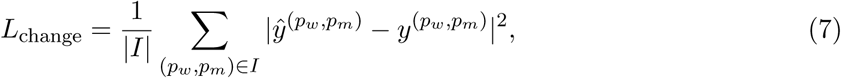

where I is the set of doublets; 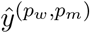 and 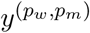 are the predicted and the true scores changes, respectively. Both scores are normalized to [0, 1] by using the min-max scaling during the optimization process.

#### Joint learning with auxiliary regressors

It has been observed in many prior works that jointly learning of multiple correlated tasks can help improve the performance of each task [4, 47]. In addition to predicting the property changes in PPI by the main multiple-layer perceptron (MLP) regressor, `MuPIPR` is able to predict the original measure of the properties for the wild-type pair and the mutant pair. This is achieved by using two auxiliary regressors, which is jointly learned to enhance the estimation of the changes. Similar to *L*_change_, we use the mean squared loss for both the wild type pair and the mutant pair: 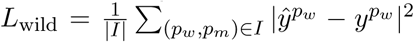, and 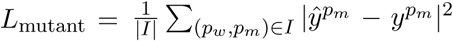. Then, the learning objective is to minimize the joint loss *L*_joint_ = *L*_change_ + *L*_wild_ + *L*_mutant_.

### 2.4 Implementation details

The BiLM for contextualized amino acid embeddings is implemented as 2 stacked layers of bidirectional LSTMs, for which the hidden dimension is set to 32. The protein sequence encoder *E*_RCNN_ consists of 3 RCNN units. The kernel size *k* of the convolution layer is set to 3 and 3-max-pooling is adopted. The hidden state of the convolution size is set to be 50, and the residual RCNN output size is set to be 100. We also zero-pad short sequences to the longest sequence length in the dataset, as a widely adopted technique in bioinformatics [37, 39, 41] for efficient training. Both the main MLP and the auxiliary MLP have 1 hidden layer with 100 neurons.

We use batched training based on the AMSGrad optimizer [45] to optimize the parameters, for which we fix the batch size as 32, the learning rate *α* as 0.005, the linear and quadratic exponential decay rates as 0.9 and 0.999 respectively. We also implement 5 simplified variants of `MuPIPR`. Specifically, `MuPIPR`-*noAux* removes the two auxiliary MLP regressors in the estimation stage. `MuPIPR`-*static* uses the same four-sequence residual RCNN encoders, but replaces the contextualized amino acid representations with the static amino acid embeddings as used in PIPR [8]. `MuPIPR`-*static-noAux* removes the two auxiliary MLP regressors and uses the static embeddings simultaneously. `MuPIPR`-*CNN* substitutes the residual RCNN encoder with convolution layers by removing the residual BiGRUs. Finally, `MuPIPR`-*CNN-noAux* further removes the auxiliary regressors. All model variants are trained until converge for each fold of the cross-validation.

### 2.5 Datasets

We obtain data from three resources for different purposes. Data from the SKEMPI (Structural database of Kinetics and Energetics of Mutant Protein Interactions) database [38] are used to conduct the task of binding affinity change estimation. The dataset for the task of BSA change estimation is constructed from the PDB (protein data bank) [5]. To pre-train the contextualized representations of amino acids, we collect protein sequences from the STRING database [48]. Details of these datasets are described below. The processed data are available at our GitHub repository.

#### SKEMPI datasets

Two datasets generated from the SKEMPI database are used for the binding affinity task. The first one is a benchmark dataset extracted by Xiong *et al.* [51]. We denote it as SKP1400m. This dataset contains the changes of binding affinity between wild-type and mutated protein complexes that are experimentally measured. These mutations include single and multiple amino acid substitutions on the protein sequences. For duplicated entries of two protein pairs with the same mutations, we take the average *∆∆G* of those reported in SKEMPI. As a result, this dataset contains 1402 doublets for 114 proteins, among which, 1131 doublets contain single-point mutations, 195 contain double-points mutations, and 76 contain three or more mutations. The second dataset is provided by Geng et al. [13], which considers only single-point mutation of dimeric proteins. It contains 1102 doublets for 57 proteins. We denote this dataset as SKP1102s. Of these 1102 doublets, the majority (759 doublets) are new entries that are not found in SKP1400m.

#### PDB dataset

We use this dataset for the task of estimating BSA changes. To construct the wild-type pairs and their mutant pairs, we extract protein sequences from PDB, and keep those with only two chains. Sequences with less than 20 amino acids are removed. Here a wild-type pair or a mutant pair refers to the two chains of a protein. We concatenate such two chains of a protein for pairwise sequence comparisons, and retain those with one amino acid substitution. This process produces 2948 doublets. To compute the true value of BSA, we first run DSSP [23] to obtain the ASA of the proteins based on the 3D structures provided by PDB. The standard estimation of BSA is calculated by taking the difference between the sum of ASA for the individual chains in a protein complex and the ASA of the protein complex [13].

#### Contextualized amino acid training corpus

We obtain the corpus to pre-train the contextualized amino acid encoder from the STRING database. A total of 66235 protein sequences of four species from the STRING database are extracted, i.e. *Homo sapiens, Bos taurus, Mus musculus* and *Escherichia coli*. These are the four most frequent species in the SKP1400m dataset.

## 3 Experiments

To demonstrate the effectiveness of `MuPIPR`, we conduct comprehensive experiments on two tasks. These two tasks estimate the mutation effects on different aspects of the alteration of the PPI properties: the *change of binding affinity (∆∆G)* and the *change of buried surface area (∆BSA)*.

### 3.1 Task 1: binding affinity change estimation

Binding affinity change estimation is a widely attempted task in previous works [6, 11, 16, 29, 43, 51]. Given a wild-type protein pair and its mutant pair, the goal is to estimate the difference of their binding affinities (*∆∆G*).

#### Evaluation protocol

Following the convention [29, 43], the performance of binding affinity change estimation is evaluated based on Pearson’s correlation coefficient (Corr) and root mean square error (RMSE), which are two widely used metrics for regression tasks. To be consistent with the baseline experiments conducted by [51] and [13], we carry out a 5-fold cross-validation on SKP1400m and a 10-fold cross-validation on SKP1102s. Note that in the we have described the configurations of `MuPIPR` in the Implementation details Section, and the study of different hyperparameter values will be presented in the Hyperparameter study Section.

#### Baseline methods

We compare `MuPIPR` with two groups of baselines for this task: (i) BeAtMusic [11], Dcomplex [33], and FoldX3 [16] are empirical linear methods; (ii) mCSM [43], BindProfX [51] and Mutabind [29] are statistical learning methods leveraging structural and/or energy features. Note that we cannot run iSEE [13] on SKP1400m, since the service of generating features used in iSEE is not provided. Hence, the comparison of iSEE is only conducted on the SKP1102s dataset used by Geng et al. [13].

#### Experimental results

The results are reported in Table 1. For *∆∆G* estimation on single-point mutation, empirical linear methods perform the worst. This shows that the pre-determined coefficients cannot be generalized well to reflect the mutation effects. On the contrary, statistical learning methods generally perform better. The best-performing baseline on SKP1400m is Mutabind, which trains a Random Forest regressor based on a variety of energetic, structural and conservation features. `MuPIPR` outperforms Mutabind by 0.038 in Corr and 0.376 kcal/mol in RMSE. As for the comparison with iSEE, we achieve Corr of 0.858 and RMSE of 1.236 kcal/mol on SKP1102s, outperforming iSEE which achieves Corr of 0.80 and RMSE of 1.41 kcal/mol. This is attributed to the fact that `MuPIPR` is able to discover important features reflecting the mutation effects.

**Table 1:** Corr and RMSE (kcal/mol) of *∆∆G* estimation by different methods using the SKP1400m dataset.

| $N_{mut}$ | All | | 1 | | 2 | | $\geq 3$ | |
| --- | --- | --- | --- | --- | --- | --- | --- | --- |
| Methods | Corr | RMSE | Corr | RMSE | Corr | RMSE | Corr | RMSE |
| BeAtMusic | - | - | 0.272 | 2.389 | - | - | - | - |
| mCSM | - | - | 0.579 | 2.019 | - | - | - | - |
| Mutabind | - | - | 0.816 | 1.675 | - | - | - | - |
| Dcomplex | 0.183 | 3.068 | 0.056 | 2.684 | -0.057 | 4.435 | 0.015 | 3.999 |
| FoldX3 | 0.480 | 2.629 | 0.470 | 2.265 | 0.289 | 3.487 | 0.095 | 4.471 |
| BindProfX | 0.690 | 1.914 | 0.663 | 1.854 | 0.584 | 2.558 | 0.840 | 2.304 |
| MuPIPR | **0.883** | **1.324** | **0.854** | **1.299** | **0.899** | **1.462** | **0.898** | **1.303** |

For cases with multiple mutations, explicit feature-based learning methods often fall short of modeling more than one mutation. Therefore, we evaluate the capabilities of `MuPIPR` in capturing the effects of different numbers of mutations (*N*_mut_) in Table 1, and compare with baselines that support multiple mutations. Dcomplex is the only empirical linear method that can model multiple mutations; however, it performs poorly in all cases. The estimations under multi-mutation cases are further impaired compared to single-point mutations. The statistical learning baseline that supports estimation upon multi-point mutations, i.e. BindProfX, shows better generalizability and offers much better performance. `MuPIPR` significantly outperforms BindProfX in all cases with different number of mutations. In particular, `MuPIPR` offers a drastic improvement of 0.280 in Corr on two-mutation cases, and that of 0.058 on cases with more than two mutations. Hence, `MuPIPR` can precisely capture the effects of multiple point mutations, where other systems typically fall short. To further demonstrate the contributions of each component of `MuPIPR`, we conduct an ablation study with simplified variants as shown in Table 2. Specifically, `MuPIPR`-*static* replaces the contextualized embedding with static amino acid embeddings; `MuPIPR`-*CNN* substitutes the residual RCNN encoder by removing the residual BiGRUs and uses convolution layers only; *noAux* remarks for removing the two auxiliary MLP regressors from the model. By adopting contextualized amino acid representations, `MuPIPR` and `MuPIPR`-*CNN* perform better than `MuPIPR`-*static* in both Corr and RMSE. This is due to the fact that contextualized representation mechanism can extract more-refined amino-acid-level features to distinguish among different contexts of the proteins. By propagating the mutation effects to surrounding amino acid representations, the subtle changes in the protein sequence can be competently captured. The complete version of `MuPIPR` outperforms `MuPIPR`-*CNN* with the improvement of 0.027 in Corr and 0.12 kcal/mol in RMSE. This shows that residual RCNN is superior in leveraging sequential and local significant features that are important to predict PPI properties.

**Table 2:** Corr and RMSE (kcal/mol) of *∆∆G* estimation by different variants of `MuPIPR` using the SKP1400m dataset.

| $N_{mut}$ | All | | 1 | | 2 | | $\geq 3$ | |
| --- | --- | --- | --- | --- | --- | --- | --- | --- |
| MuPIPR variants | Corr | RMSE | Corr | RMSE | Corr | RMSE | Corr | RMSE |
| MuPIPR-*static-noAux* | 0.773 | 1.832 | 0.741 | 1.783 | 0.794 | 2.032 | 0.744 | 2.001 |
| MuPIPR-*static* | 0.795 | 1.721 | 0.754 | 1.684 | 0.821 | 1.946 | 0.858 | 1.654 |
| MuPIPR-*CNN-noAux* | 0.819 | 1.596 | 0.779 | 1.559 | 0.829 | 1.728 | 0.797 | 1.773 |
| MuPIPR-*CNN* | 0.856 | 1.449 | 0.822 | 1.410 | 0.869 | 1.663 | 0.867 | 1.435 |
| MuPIPR-*noAux* | 0.853 | 1.462 | 0.829 | 1.406 | 0.848 | 1.751 | 0.860 | 1.473 |
| MuPIPR | **0.883** | **1.324** | **0.854** | **1.299** | **0.899** | **1.462** | **0.898** | **1.303** |

To verify the effectiveness of adopting auxiliary regressors, we train the *noAux* variants to learn *∆∆G* without the original binding affinities, *∆G*_w_ for wild-type and *∆G*_m_ for mutant. Table 2 demonstrates that the variants with auxiliary regressors (`MuPIPR`-*static*, `MuPIPR`-*CNN* and `MuPIPR`) consistently outperform their counterparts (`MuPIPR`-*static-noAux*, `MuPIPR`-*CNN-noAux* and `MuPIPR`-*noAux*) by 0.022, 0.037 and 0.030 in Corr, respectively. The auxiliary MLP regressors jointly learn the original scores, which can effectively guide the learning of *∆∆G* and hence improve the performance. We further evaluate the performance of the auxiliary MLP regressors in estimating the original binding affinities. Table 4 shows that while all variants perform comparably well on estimating the *∆G*_*w*_, `MuPIPR`-*CNN* and `MuPIPR` ameliorate the estimation of *∆G_m_* through contextualized amino acid representations. To remove the bias of training and testing of our model on proteins with high sequence similarity, we evaluate the performance of `MuPIPR` by removing homologous proteins at different sequence similarity thresholds. Specifically, we use CD-HIT [12, 31] to cluster the wild-type protein sequences from SKP1102s with 75% and 45% as thresholds. For each cluster, we select one protein as the representative and remove the entries of other proteins. Accordingly, we retain 652 and 637 protein pairs at the thresholds of 75% and 45% respectively. After conducting 10-fold cross-validation, `MuPIPR` achieves a Corr of 0.794 and an RMSE of 1.340 kcal/mol on the subset with less than 75% sequence identity. On the subset with less than 45% sequence identity, it achieves a Corr of 0.758 and an RMSE of 1.434 kcal/mol. By removing the homologous proteins in the data, `MuPIPR` still performs reasonably well based on the evaluation metrics. This demonstrates `MuPIPR` is robust in identifying the important physicochemical changes regarding the changes of binding affinity.

#### Blind test evaluations

To show the generalizability of `MuPIPR`, we evaluate the performance of different methods on two external validation sets. The first blind test is provided by Benedix et al. [3], which contains the mutations on chain B of the interleukin-4 receptor (PDB ID: 1IAR). We denote this set as the NM test set. The second blind test contains the mutations of the MDM2-p53 complex (PDB ID: 1YCR) provided by Geng et al. [13]. These mutations have been shown important in cancer development [50].

We follow the same experimental settings as Geng et al. [13] and apply the same data filtering criteria for the test sets. In addition, we use BLASTp [2] to verify that these test sets are unknown to the training phase. We search all the sequences from the test sets against all the wild type sequences from SKP1102s and find that the best E-values are 0.35, 2.1, and 0.15 for the interleukin-4 receptor, MDM2, and p53, respectively. The BLASTp results indicate that the sequences of these test sets are significantly different from those in the training data.

We compare `MuPIPR` with four baselines including mCSM, BeAtMuSic, FoldX and iSEE. The results reported in Table 3 show that `MuPIPR` outperforms other methods on NM by achieving a 0.74 in Corr and a 1.13 kcal/mol in RMSE. While all baseline methods fall short of predicting *∆∆G* on MDM2-p53, `MuPIPR` drastically outperforms others on MDM2-p53 with a significantly higher Corr of 0.43 and a lower RMSE of 0.70 kcal/mol. The comparative results of `MuPIPR` on the NM and MDM2-p53 sets demonstrate that `MuPIPR` is not over-fitting and can be generalized to mutations on complexes with very limited sequence similarity in the training set.

**Table 3:**
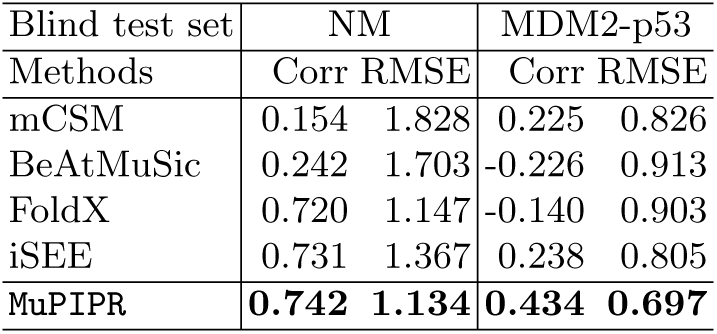
Corr and RMSE (kcal/mol) of *∆∆G* estimation on the external validation sets NM and MDM2-p53.

**Table 4:** Corr and RMSE (kcal/mol) of binding affinity estimation by different variants of `MuPIPR` using the SKP1400m dataset.

| | $\Delta G_w$ | | $\Delta G_m$ | |
| --- | --- | --- | --- | --- |
| Methods | Corr | RMSE | Corr | RMSE |
| MuPIPR-*static* | **0.978** | **0.722** | 0.860 | 1.775 |
| MuPIPR-*CNN* | 0.964 | 0.872 | 0.901 | 1.520 |
| MuPIPR | 0.966 | 0.872 | **0.903** | **1.483** |

#### Analysis of mutation types

Consider that different types of point mutations can occur to a protein sequence and yield different impacts on the protein complex, we provide an analysis to demonstrate how `MuPIPR` captures the mutation effect for specific types of mutations. In particular, we consider the most common point mutation where one amino acid is mutated to alanine. For simplicity, we regard such a mutation as an “ALA” mutation, and others as a “NonALA” mutation. In Figure 3, we use the scatter plot to show the correlations between predicted and experimental *∆∆G* values for these two groups in SKP1102s, in which 335 out of 1102 samples are ALA mutations and 767 are NonALA mutations. `MuPIPR` achieves a Corr of 0.61 and an RMSE of 1.17 kcal/mol for the ALA mutations, and a Corr of 0.89 and an RMSE of 1.24 kcal/mol for the NonALA mutations. We compare our findings to the results reported by iSEE [13]. iSEE attains lower Corrs (0.50 for ALA and 0.84 for NonALA) and higher RMSEs (1.21 kcal/mol for ALA and 1.48 kcal/mol for NonALA) than `MuPIPR`. Thus, `MuPIPR` is more effective in capturing the effects of alanine mutations.

**Fig. 3:**
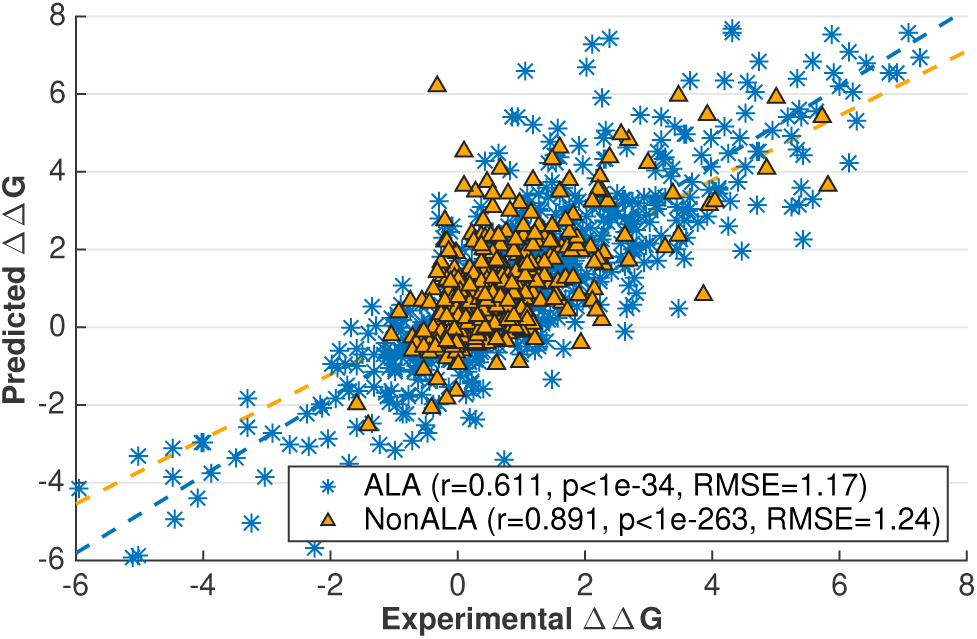
Correlations between predicted and experimental *∆∆G* values for different types of mutated amino acids (i.e. “ALA” and “NonALA”) in the SKP1102s dataset.

We further investigate the ability of `MuPIPR` on predicting the mutation effects of all 20 amino acids. The results and detailed analysis are presented in Supplemental Materials. Figure S2 summarizes the distribution of prediction errors for different amino acids. Overall, `MuPIPR` performs consistently well on different mutation types, demonstrating that the predictions from `MuPIPR` are not biased toward any specific amino acid.

### 3.2 Task 2: Buried surface area change estimation

The second task estimates the change of buried surface area (*∆*BSA) of two chains of a protein complex upon mutations.

#### Evaluation protocol

Same as the first task, the performance of BSA change estimation is evaluated with Corr and RMSE. We also carry out a 5-fold cross-validation on the PDB dataset.

#### Baseline method

To the best of our knowledge, the state-of-the-art methods only focus on estimating the solvent accessible surface area (ASA), instead of BSA and *∆*BSA. Among these methods, we choose the best performing one, NetSurfP-2.0 [25], as our baseline. NetSurfP-2.0 is a deep learning model, which learns an architecture composed of convolutional and LSTM networks using sequence information. It estimates the ASA at the residue level for each protein sequence. The ASA of a chain of protein is then calculated by summing up the ASA of all residues in the sequence. The BSA and its change are then calculated based on the predicted ASA. In addition, we take three variants of `MuPIPR` into comparison, including `MuPIPR`-*static*, `MuPIPR`-*noAux* and `MuPIPR`-*CNN* as described in Task 1.

#### Experimental results

`MuPIPR` provides a direct estimation of *∆*BSA, BSA_*w*_ for wild-type and BSA_*m*_ for mutant, without pre-estimating the ASA. Table 5 shows the estimations on the PDB dataset. All the `MuPIPR` variants drastically outperform NetSurfP-2.0 on both BSA and *∆*BSA. The worst-performing variant `MuPIPR`-*static*, has already outperformed NetSurfP-2.0 with more than 1.4-fold of Corr on both BSA_*w*_ and BSA_*m*_, As for *∆*BSA estimation, `MuPIPR`-*static* almost doubles (1.9-fold) the Corr reported by NetSurfP-2.0. Similar to what we have observed in Task 1, incorporating both contextualized amino acid representations and auxiliary regressors leads to the best performance. `MuPIPR` offers a fold change of 2.1 in Corr over the results of NetSurfP-2.0. This once again indicates the importance of contextualized representations in highlighting the mutations in sequences, and the benefits of jointly capturing the original BSA.

**Table 5:** Corr and RMSE (10^2^*Å*^2^) of BSA and *∆*BSA estimation by different methods using the PDB dataset.

| | $BSA_w$ | | $BSA_m$ | | $\Delta BSA$ | |
| --- | --- | --- | --- | --- | --- | --- |
| Method | Corr | RMSE | Corr | RMSE | Corr | RMSE |
| NetSurfP-2.0 | 0.670 | 25.994 | 0.685 | 22.731 | 0.329 | 3.405 |
| MuPIPR-*static* | 0.978 | 4.833 | 0.980 | 4.681 | 0.624 | 2.031 |
| MuPIPR-*noAux* | NA | NA | NA | NA | 0.667 | 2.021 |
| MuPIPR-*CNN* | 0.985 | 4.079 | 0.987 | 3.794 | 0.673 | 1.913 |
| MuPIPR | **0.986** | **3.844** | **0.988** | **3.616** | **0.695** | **1.859** |

### 3.3 Case studies of contextualized amino acid embeddings

We further demonstrate the effectiveness of contextualized representations in capturing the effect of mutations on both tasks using specific examples.

Table 6 depicts 3 different cases of mutations on the barnase-barstar complex (1B3S): (i) The mutation at position 25 of Chain D (*Phe* to *Tyr*) leads to a small change of binding affinity; (ii) The mutation at position 102 of Chain A (*Ala* to *His*) causes a substantial change; (iii) Combining the mutations of these two brings a significant change in the binding affinity of this complex. For each case, we also investigate the *L*_2_ differences of sequence pair encodings between the wild-type pair and the mutant pair. The corresponding *L*_2_ differences under static and contextualized amino acid representations are denoted by *D*_static_ and *D*_*Eco*_, respectively. *D*_static_ is consistently small, which indicates the subtle changes to sequences to be inadequately captured. As a result, its *∆∆G* is more erroneously estimated. On the other hand, the *∆∆G* for *D*_*Eco*_ is estimated more accurately. This shows the contextualized representations amplify the mutation effects, as reflected by *D*_*Eco*_. These observations show that the contextualized amino acid embeddings better benefit the downstream sequence encoder to capture the subtle changes of a sequence, such as point mutations, and lead to a more accurate estimation of binding affinity changes.

**Table 6:** Case study of the effects of contextual amino acid embeddings on two chains of a protein complex, 1B3S A for barnase and 1B3S D for barstar.

| Mutation(s) | $L_2$ distances | | Ground truth<br>$\Delta\Delta G$ | $\Delta\Delta G$ Predictions | |
| --- | --- | --- | --- | --- | --- |
| | $D_{static}$ | $D_{E_{co}}$ | | MuPIPR-*static* | MuPIPR |
| D_F30Y | -0.0027 | -0.0073 | 0.596 | -5.534 | 0.185 |
| A_A102H | 0.0021 | -0.0518 | -5.682 | -5.032 | -6.141 |
| A_A102H, D_F30Y | -0.0006 | -0.0591 | -9.549 | -2.082 | -8.868 |

For the change of BSA estimation, Figure 4 depicts the 3D structures of two chains of the Human Insulin complex. *Phe* at position 25 of Chain B is mutated to *His*, causing a significant conformational alteration of the complex. The BSA therefore changes from −1401*Å*^2^ to −1055*Å*^2^. The true *∆*BSA here (−346*Å*^2^) is more precisely estimated by `MuPIPR` with an absolute error of 1.61 than `MuPIPR`-*static* with an absolute error of 126.95. The static amino acid embeddings again fall short of capturing the changes upon mutation and lead to a notably less accurate estimation. In addition, we provide a comprehensive analysis of all the samples in the PDB dataset to further demonstrate the merits of using the contextualized amino acid embeddings over the static embeddings in Supplemental Materials Figure S3.

**Fig. 4:**
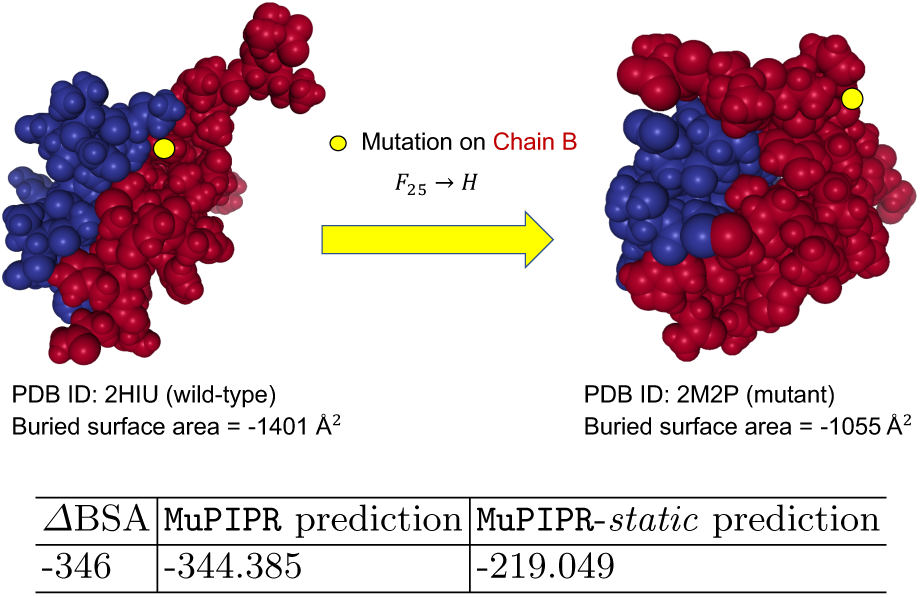
Mutation effects on structures and BSA. The structures of Chain A and Chain B of the Human Insulin protein complex are depicted respectively in blue and red. The mutation is highlighted in yellow. The wild-type (2HIU) and mutant (2M2P) complexes are retrieved from PDB.

### 3.4 Hyperparameter study

We conduct the study on the configuration of critical factors that affects the performance of the amino acid contextualized embeddings: a) the dimensionality of hidden states for *E*_*co*_; b) the number of LSTM layers used in *E*_*co*_; c) the dimensionality of hidden states for the residual RCNN; d) the number of residual RCNN units. Fig 5 shows the performances of different settings for these 4 hyperparameters based on the change of binding affinity estimation task using the SKP1400m dataset.

**Fig. 5:**
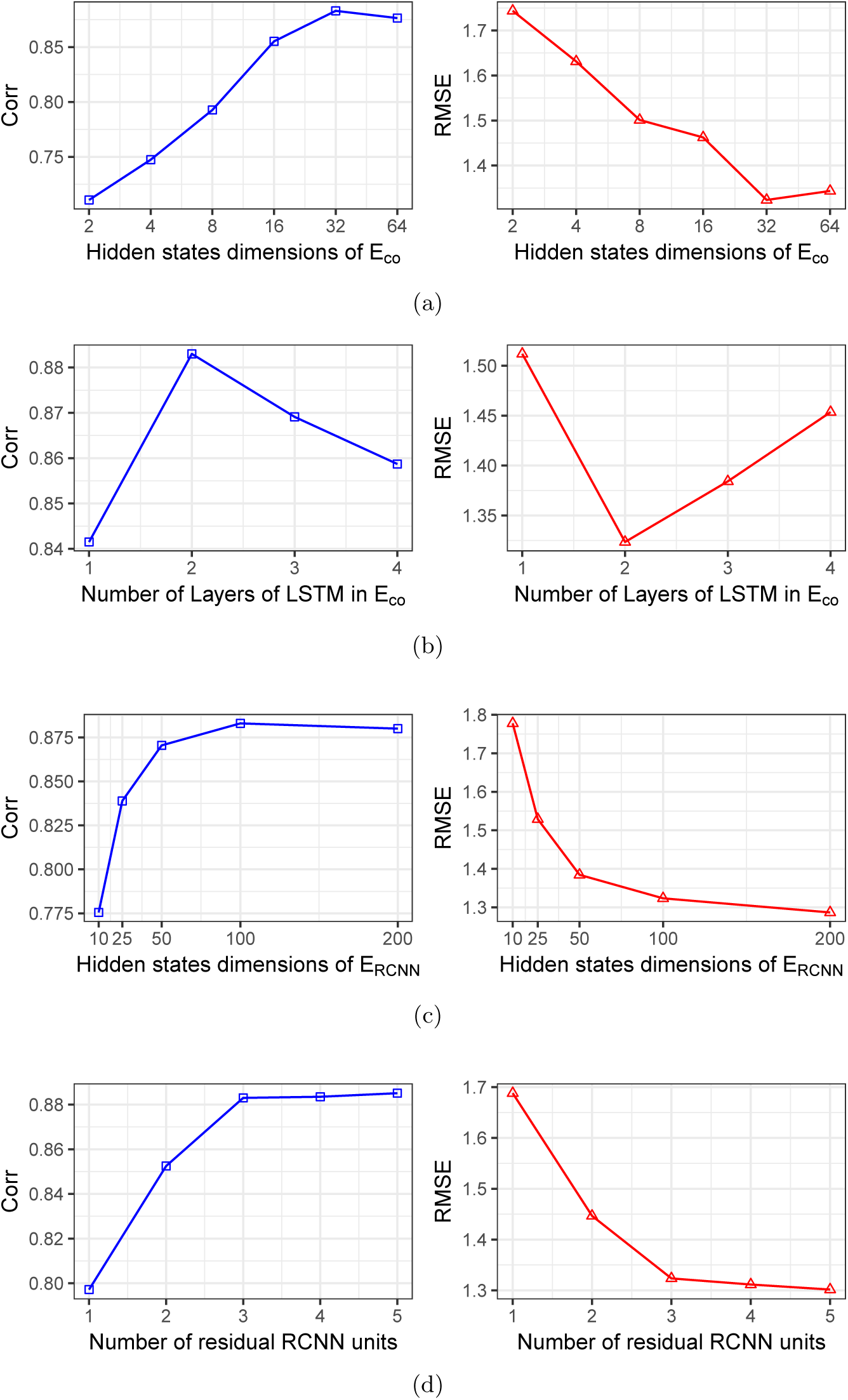
Performance evaluation on using different hyperparameters on the SKP1400m dataset for *∆∆G* estimation. The Pearson’s correlation coefficient (Corr) and the root mean square error (RMSE) are reported in blue squares (left) and red triangles (right), respectively.

The dimensions of hidden states for *E*_*co*_ are chosen from {2, 4, 8, 16, 32, 64}. As illustrated in Fig 5a, the performance of `MuPIPR` gets better as we increase the dimensionality of the hidden states until it reaches 32 and then the performance slightly drops with dimension of 64. The number of LSTM layers in *E*_*co*_ is another factor. Different layers of the stacked LSTM captures different widths of the neighbouring contexts of the amino acid on the sequence and we examine the cases with 1 to 4 layers. Fig 5b shows that 2 layers is the optimum.

Apart from the parameters of *E*_*co*_, we also exam the hidden states and the number of residual RCNN units for *E*_*RCNN*_. As shown in Fig 5c, the model offers the best performance when the hidden states of residual RCNN is 100. As for the residual RCNN unit, it contributes to the different levels of granularity in feature aggregation where more units correspond to more aggregation. Fig 5d demonstrates that the performances increase with more residual RCNN units. However, the improvement from 3 to 5 units is marginal because too many residual RCNN units can lead to over-compressing the features. Our framework is robust to this setting as long as there are more than 3 residual RCNN units.

## 4 Conclusion and Future Work

In this paper, we introduce a novel and comprehensive learning framework to estimate mutation effects on protein-protein interactions. Our proposed framework, `MuPIPR`, incorporates a contextualized representation mechanism of amino acids, which automatically extracts amino-acid-level features that are differentiated among different contexts of the proteins. Therefore this mechanism propagates the mutation effects to surrounding amino acid representations. By incorporating the contextualized representation mechanism to a carefully designed 4-fold Siamese deep learning architecture, `MuPIPR` effectively captures the PPI property changes between a wild-type pair and a mutant pair of proteins. Moreover, auxiliary regressors are provided to further improve estimation whenever original measures of the PPI property are available. Extensive experiments conducted on two different tasks demonstrate the promising performance of `MuPIPR`. As a future direction, we plan to explore more tasks of PPI property changes upon mutations. To further improve the performance, we seek to incorporate multi-task learning [47] to augment the learning of tasks that complement each other.

## Supplementary Materials

### I Analysis on SKEMPI version 2

We evaluate `MuPIPR` and three other predictors (FoldX, mCSM, and iSEE) on a blind set (SKPv2-487 dataset) constructed from the recently released SKEMPI 2.0 [22]. This dataset contains 487 mutations in 56 protein complexes that have not been seen in the training set. `MuPIPR` is trained on SKEMPI v1 (SKP1102s). Notably, none of the predictors perform well as shown in Table S1. FoldX performs the best in Corr (0.34) but the worst in RMSE (1.53 kcal/mol). mCSM, iSEE, and `MuPIPR` all achieve a Corr of 0.25. Among them, iSEE presents a slightly better RMSE of 1.32 kcal/mol.

Using the SKEMPI v1 as the learning resource to predict new proteins in SKEMPI v2 is very challenging. `MuPIPR` extracts the protein features from scratch on raw sequences, which inevitably fall short for cases where test data have distinct distributions on the sequence level. Therefore, `MuPIPR` is mostly suitable for estimating the mutation effects when there is adequate sequence relatedness. To support our assumption, we examine the relationship between RMSE and sequence similarity. Specifically, we run BLASTp for all wild-type sequences in SKPv2-487 (test set) against the SKP1102s dataset (training set) and record the smallest E-value for each test sequence from the BLASTp results. Then, We divide the protein pairs in the test set into four groups based on their E-values and report the RMSE of all methods as shown in Figure S1. The smaller E-value reflects a higher sequence similarity between the test and training. It is a clear trend that with the increasing of the E-value, the prediction performance drops more significantly. This demonstrates the importance of adequate sequence relatedness between the training and test set, and shows an intrinsic limitation of `MuPIPR`. Yet, we note that `MuPIPR` still offers a close performance to the other methods that employ experimental features.

**Table S1:** Corr and RMSE (kcal/mol) of binding affinity estimation by different methods on the SKPv2-487 dataset.

| Methods | Corr | RMSE |
| --- | --- | --- |
| mCSM | 0.25 | 1.35 |
| iSEE | 0.25 | 1.32 |
| FoldX | 0.34 | 1.53 |
| MuPIPR-CNN | 0.25 | 1.36 |

### II Prediction error by mutation types

We break down the performance of `MuPIPR` by all 20 amino acid mutation types based on the experiments conducted on the SKP1400m dataset. Figure S2 summarizes the distribution of prediction errors for different amino acids. These amino acids are regarded as the products of the mutation. The prediction error is calculated as the difference between experimental and predicted *∆∆G* values. The boxplots show that `MuPIPR` consistently maintains high-quality predictions with all mutation types. This demonstrates that the predictions from `MuPIPR` are not biased toward specific mutation types.

### III Analysis on contextualized amino acid embeddings

**Fig. S1:**
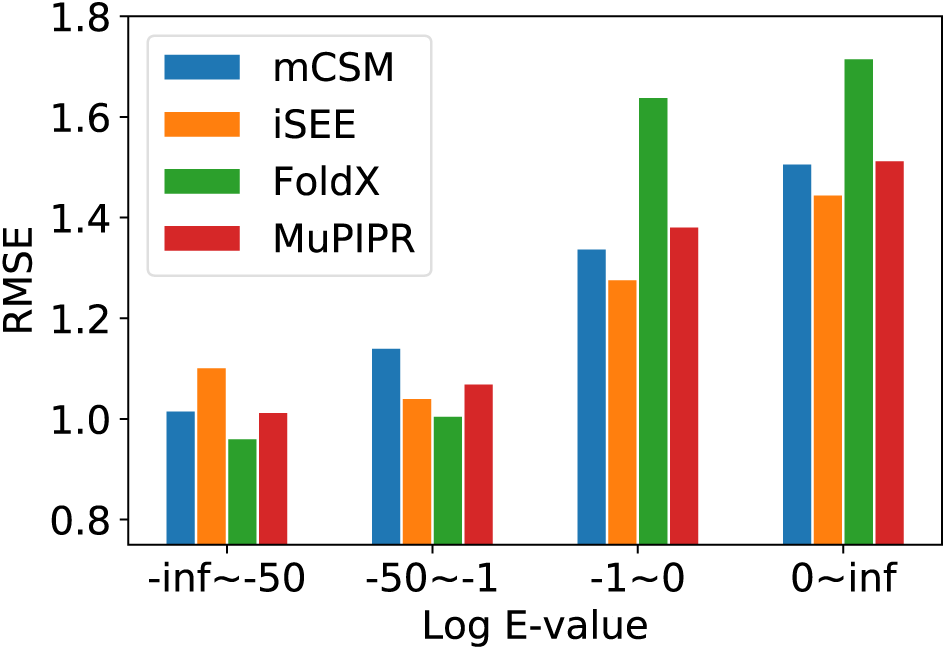
The performance of different predictors on SKEMPI v2 based upon different thresholds of sequence similarity (by the log E-value) when compared with the SKEMPI v1 training set. The number of samples in the four bins are 128, 78, 104, and 177, respectively.

**Fig. S2:**
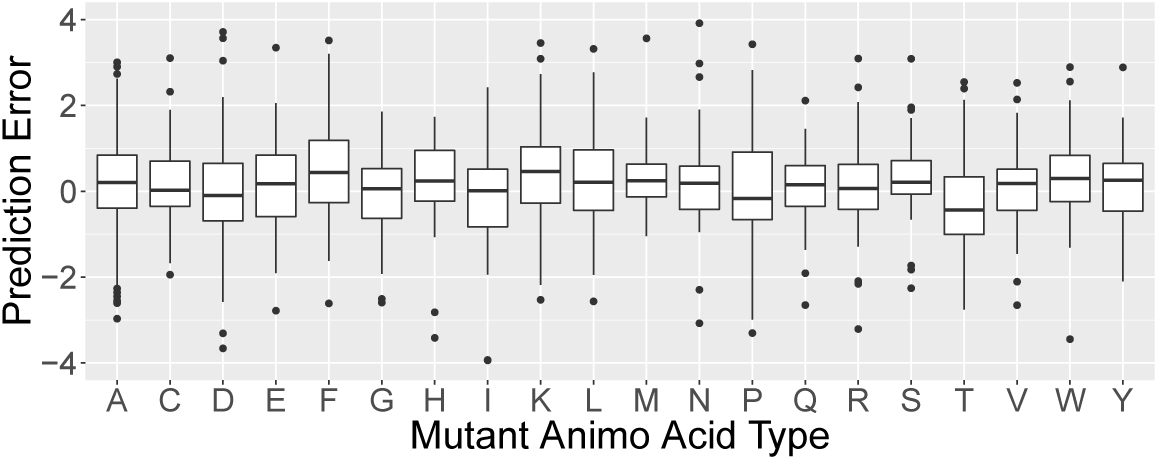
Boxplots of prediction errors for different mutant types from the SKP1400m dataset.

**Fig. S3:**
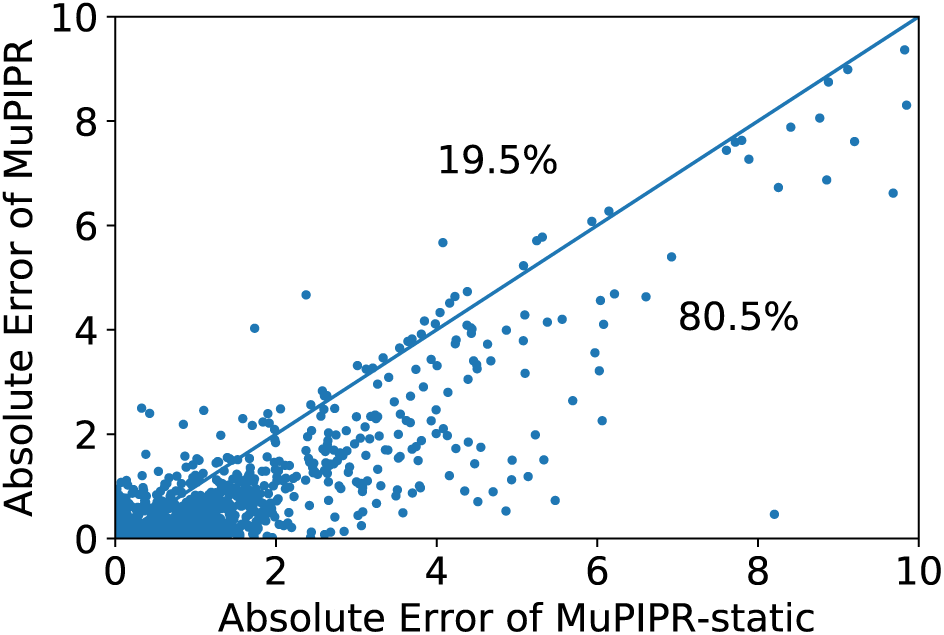
Scatter plot to compare the absolute errors by the complete version of `MuPIPR` and `MuPIPR`-static variant. The majority of points (*∼* 80.5%) are below the diagonal line, showing the predictions by the complete model to be generally more accurate.

We present a comparative study on `MuPIPR` with and without contextualized amino acid embeddings based on the change of BSA estimation task. Based on the predictions, we calculate the absolute errors for both model variants and use a scatter plot to compare this metric in Figure S3. The absolute error is calculated as the absolute difference between experimental and predicted *∆∆BSA* values. As we can see, there are many more points (*∼* 80.5%) below the diagonal line than above (*∼* 19.5%), which indicate that most cases predicted by the complete version of `MuPIPR` have smaller errors than those predicted by `MuPIPR`-static. The errors of the complete version of `MuPIPR` are also statistically significantly smaller than the errors of `MuPIPR`-static by the one-tailed paired Student’s t-test (t-statistic of −5.58 and *p*-value *<* 0.001). This improvement in performance is due to the contribution of the contextualized amino acid embeddings, which help the downstream sequence encoder to capture the subtle changes of a sequence, therefore leading to a more accurate estimation of BSA changes.

## Notes

https://github.com/guangyu-zhou/MuPIPR

